# *pDriver* : A novel method for unravelling personalised coding and miRNA cancer drivers

**DOI:** 10.1101/2020.04.23.058727

**Authors:** Vu VH Pham, Lin Liu, Cameron P Bracken, Thin Nguyen, Gregory J Goodall, Jiuyong Li, Thuc D Le

**Affiliations:** UniSA STEM, University of South Australia, Mawson Lakes, SA 5095, Australia; Centre for Cancer Biology, an alliance of SA Pathology and University of South Australia, Adelaide, SA 5000, Australia; Department of Medicine, The University of Adelaide, Adelaide, SA 5005, Australia; Applied Artificial Intelligence Institute, Deakin University, Australia

## Abstract

**Motivation:** Unravelling cancer driver genes is important in cancer research. Although computational methods have been developed to identify cancer drivers, most of them detect cancer drivers at population level. However, two patients who have the same cancer type and receive the same treatment may have different outcomes because each patient has a different genome and their disease might be driven by different driver genes. Therefore new methods are being developed for discovering cancer drivers at individual level, but existing personalised methods only focus on coding drivers while microRNAs (miRNAs) have been shown to drive cancer progression as well. Thus, novel methods are required to discover both coding and miRNA cancer drivers at individual level.

**Results:** We propose the novel method, *pDriver*, to discover personalised cancer drivers. *pDriver* includes two stages: (1) Constructing gene networks for each cancer patient and (2) Discovering cancer drivers for each patient based on the constructed gene networks. To demonstrate the effectiveness of *pDriver*, we have applied it to five TCGA cancer datasets and compared it with the state-of-the-art methods. The result indicates that *pDriver* is more effective than other methods. Furthermore, *pDriver* can also detect miRNA cancer drivers and most of them have been confirmed to be associated with cancer by literature. We further analyse the predicted personalised drivers for breast cancer patients and the result shows that they are significantly enriched in many GO processes and KEGG pathways involved in breast cancer.

**Availability and implementation:** *pDriver* is available at https://github.com/pvvhoang/pDriver

**Supplementary information:** Supplementary data are available at *Bioinformatics* online.

## 1 Introduction

Cancer driver genes play a vital role in cancer initialisation and development. Unravelling cancer drivers and their regulatory mechanisms is critical to the understanding of cancer and the design of effective cancer treatments. Many computational methods have been developed to identify cancer drivers, mainly including mutation-based methods and network-based methods. Mutation-based methods discover cancer drivers by investigating the characteristics of mutations. For instance, MutSigCV (Lawrence *et al.*, 2013) evaluates the significance of mutations in genes, OncodriveFM (Gonzalez-Perez and Lopez-Bigas, 2012) and DriverML (Han *et al.*, 2019) examine the functional impact of mutations, OncodriveCLUST (Tamborero *et al.*, 2013) uses recurrence of mutations, ActiveDriver (Reimand and Bader, 2013) looks at enrichment in externally defined regions, and CoMEt (Leiserson *et al.*, 2015) uses mutual exclusivity. Network-based methods detect cancer drivers by evaluating the role of genes in biological networks like DriverNet (Bashashati *et al.*, 2012), MEMo (Ciriello *et al.*, 2012), HotNet (Reyna *et al.*, 2018), NetSig (Horn *et al.*, 2018), and CBNA (Pham *et al.*, 2019). All these methods detect cancer drivers at the population level. However, different patients have different genomes and their diseases may be driven by different genes, and therefore, two patients having the same cancer type and receiving the same treatment may have different outcomes. Thus, there is a need to study cancer drivers specific to individual patients, called personalised cancer drivers in this paper.

Recently, some methods have been developed to identify personalised cancer drivers such as DawnRank (Hou and Ma, 2014), SCS (Guo *et al.*, 2018), and PNC (Guo *et al.*, 2019). DawnRank considers mutated genes with higher connectivity in the gene regulatory network are more impactful and identifies such genes by applying PageRank (Page *et al.*, 1998; Brin and Page, 1998) to the gene network. To evaluate the influence of genes, DawnRank uses each individual patient’s gene expression data but the same gene network is used for all patients. On the other hand, SCS builds a gene network for each patient using the patient’s gene expression data and the gene expression data of the neighbour (i.e. a normal sample). SCS identifies cancer drivers of a patient as the minimal set of mutated genes which controls the maximum number of differentially expressed genes in that patient’s network. Similarly, PNC uses the gene expression data of a tumour and its neighbour to build a personalised network. However, PNC only keeps edges which have co-expression p-value of the two nodes less than 0.05 in one state and greater than 0.05 in the other state. Then it converts the gene network to a bipartite graph. In the bipartite graph, the nodes on the top represent genes and the nodes on the bottom represent edges. The cancer drivers predicted by PNC are the minimum set of genes on the top which covers all the edges on the bottom in the bipartite graph.

Although these methods can be used to identify personalised cancer driver genes, they have their own limitations. DawnRank uses the same gene regulatory network for all patients and ignores the network information which is specific to each patient (Hou and Ma, 2014). Although SCS and PNC integrate the genetic data of each patient to build personalised gene networks, they both require the gene expression data of a tumour and its normal neighbour. However, identifying the neighbour of a tumour is not easy and in some cases, the normal neighbour does not exist. Furthermore, these methods only uncover coding driver genes while cancer drivers might be non-coding genes (e.g. miRNAs). Because protein-coding regions only account for about two percent of the human genome (Yang *et al.*, 2016a), a large percentage of mutations might exist in non-coding regions, and thus non-coding genes may act as cancer drivers too (Puente *et al.*, 2015; Weinhold *et al.*, 2014). Consequently, novel and effective methods are needed for both coding and non-coding personalised cancer drivers.

In this paper, we develop a novel method, *pDriver*, to discover personalised coding and miRNA cancer drivers. *pDriver* provides a thorough “treatment” to the problems with existing methods as it has been designed with personalised gene regulation in mind and has considered gene regulation network as a control system whose functioning is driven by some key components. This view has led us to adopt and combine innovatively the techniques from different disciplines to develop *pDriver*. Similar to the existing methods like SCS and PNC, *pDriver* also takes the advantages of gene regulatory networks and constructs the gene regulatory network for each patient but unlike SCS and PNC, we do not require the gene expression data of the neighbour of a patient, which greatly enhances the usability of *pDriver* since in practice it is often difficult to find a matching “neighbour” for a patient. Furthermore, *pDriver* provides a more comprehensive coverage for personalised driver discovery, by considering both coding and non-coding drivers.

In particular, to make use of gene network information specific to a patient, we firstly build gene regulatory networks for each cancer patient based on the matched mRNAs, Transcription Factors (TFs), and miRNAs expression data of the patient. Applying LIONESS, a method to estimate sample-specific regulatory networks (Kuijjer *et al.*, 2019), we build the gene network for a patient based on the difference between the network built from the data of all patients and the network built from the data of all patients except the patient under consideration. Then based on the directed PPI network (Vinayagam *et al.*, 2011) and the existing gene interaction databases we refine each patient’s gene network by removing interactions not supported by the PPIs and the gene interaction databases. We further remove edges with a low weight to keep only edges which have a strong connection for each patient. To predict cancer drivers of a patient, we need to identify genes which play the critical role in controlling the whole gene network of the patient. System controllability has been a central topic studied for decades, especially in the engineering discipline. However, it is computationally prohibitive to apply classic control theory, such as Kalman’s controllability theory (Kalman, 1963) to a gene regulation network due to its high complexity. Thanks to the recent work by Liu *et al.* (2011), which has provided an analytical method to identify the set of driver nodes in a complex system modelled by a weighted directed network. Following the Network Control method in Liu *et al.* (2011), in a system, there exists a set of nodes which are critical to the control of the working of the system such that removing such a node will require more nodes to control the network, which nicely mimics the role of a cancer driver in a gene regulatory network. Thus we adopt this Network Control method to identify cancer drivers as such critical nodes. As we will show in Section 3.5, comparing to other methods for finding influential nodes in a network, the Network Control method outperforms the other methods since it has captured the control mechanism of gene regulation while the others discover influential nodes only from the information propagation perspective.

We apply *pDriver* to five TCGA cancer datasets and validate the results with the Cancer Gene Census (CGC), in comparison with the state-of-the-art cancer driver prediction methods, including 3 personalised driver prediction methods (DawnRank (Hou and Ma, 2014), PNC (Guo *et al.*, 2019), and SCS (Guo *et al.*, 2018)) and 5 population level driver prediction methods (ActiveDriver (Reimand and Bader, 2013), DriverML (Han *et al.*, 2019), DriverNet (Bashashati *et al.*, 2012), MutSigCV (Lawrence *et al.*, 2013), and OncodriveFM (Gonzalez-Perez and Lopez-Bigas, 2012)). Since there is no ground truth available for individual patients’ cancer drivers, following the same approach in existing literature on personalised driver prediction, we aggregate the results of personalised coding drivers discovered by *pDriver* (and the other 3 personalised driver prediction methods) for the comparison. In validating the miRNA cancer drivers discovered by *pDriver*, we use OncomiR (Wong *et al.*, 2018), a database of miRNA dysregulation in pan-cancer. Over 50% of the discovered miRNA drivers are in OncomiR and several predicted miRNA drivers are confirmed to be involved in different cancer types by literature.

In addition, the personalised drivers found by *pDriver* are significantly enriched in various GO biological processes and KEGG pathways related to cancer. We focus on personalised rare coding drivers which may be drivers of many patients but they have a low mutation frequency, as these drivers are usually missed by other cancer driver identification methods, especially methods based on gene mutations. The analysis of the distribution of the personalised rare coding drivers found by *pDriver* in breast cancer subtypes reveals that Basal cancer patients are driven by different rare coding drivers while other subtypes such as Luminal A cancer patients are usually driven by the same rare coding drivers. This finding may be associated with the great heterogeneity of Basal breast cancer.

## 2 Datasets and methods

### 2.1 Datasets

In this paper, we use five TCGA datasets (The Cancer Genome Atlas Research *et al.*, 2013): breast invasive carcinoma (BRCA), lung adenocarcinoma (LUAD), lung squamous cell carcinoma (LUSC), kidney renal clear cell carcinoma (KIRC), head and neck squamous cell carcinoma (HNSC).

These datasets contain the gene expression data of cancer patients and they are used to construct gene networks. The TF list is obtained from Lizio *et al.* (2017) and we use this list to distinguish TF genes from other coding genes. Several gene interaction datasets are used to refine the constructed gene networks, including PPIs (Vinayagam *et al.*, 2011), TransmiR 2.0 (Wang *et al.*, 2010), TargetScan 7.0 (Agarwal *et al.*, 2015), miRTarBase 6.1 (Chou *et al.*, 2016), TarBase 7.0 (Vlachos *et al.*, 2015), and miRWalk 2.0 (Dweep and Gretz, 2015). The details of these datasets will be discussed in the sections below and they are available at https://github.com/pvvhoang/pDriver.

### 2.2 pDriver

#### 2.2.1 The workflow of *pDriver*

As shown in Fig. 1, *pDriver* contains two stages to identify personalised coding and miRNA cancer drivers: (1) Constructing the miRNA-TF-mRNA network for each cancer patient, and (2) Identifying coding and miRNA cancer drivers for each patient based on the personalised network built in the first stage. The details of the two stages are described below.

**Fig. 1.**
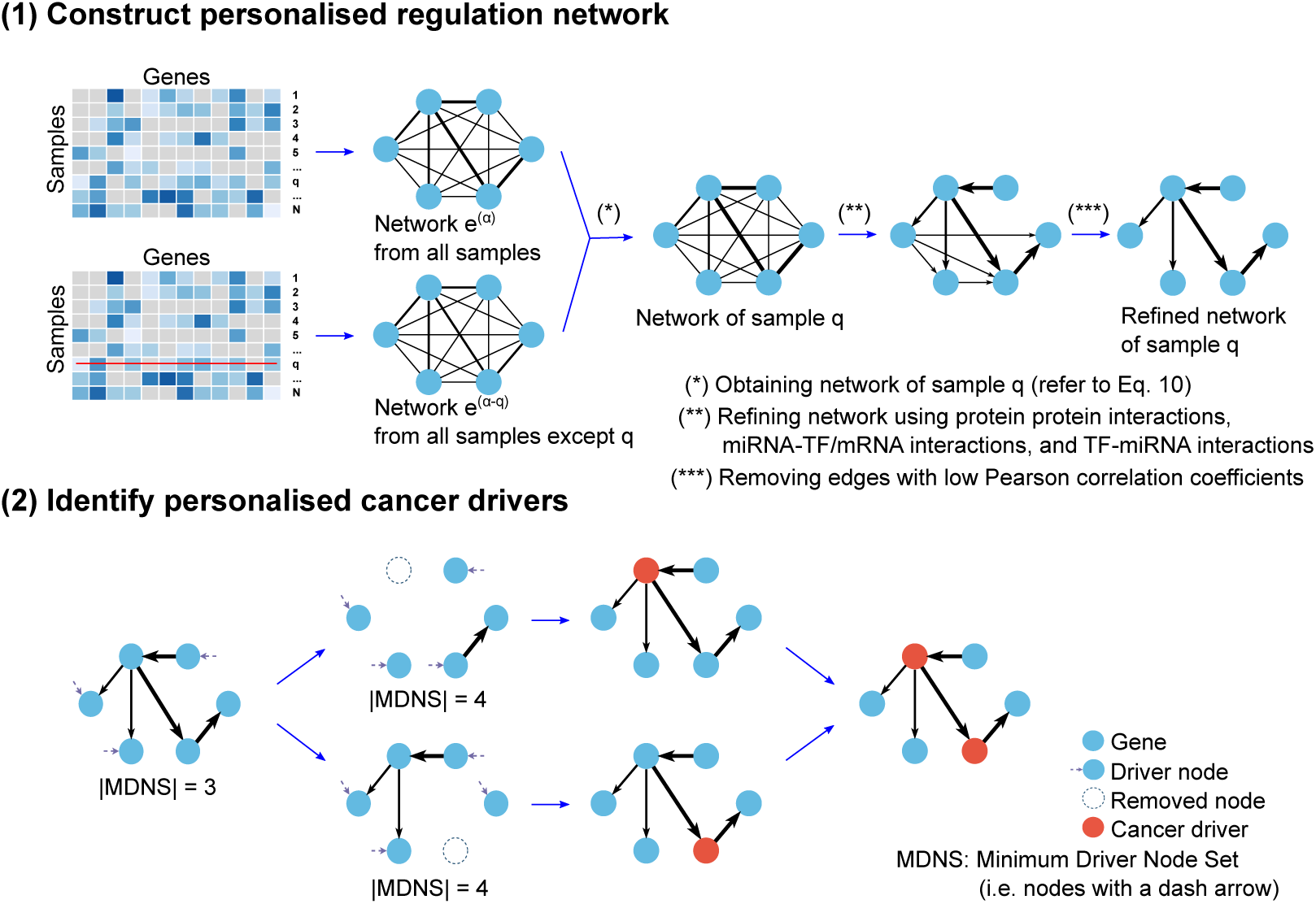
An illustration of *pDriver*. (1) Building the gene network for each cancer patient based on gene expression data and refine these patient-specific networks using existing gene interaction databases (including protein protein interactions, miRNA-TF/mRNA interactions, and TF-miRNA interactions) to remove unreal interactions and using Pearson correlation coefficients between genes to keep only edges which have a strong correlation in each patient, and (2) Identifying coding and miRNA cancer drivers for each patient by evaluating the role of genes in the personalised network.

##### Stage (1) Constructing personalised miRNA-TF-mRNA networks

**Step 1.** Prepare gene expression data. We obtain the expression data of coding genes and miRNAs of matched samples from the TCGA datasets (BRCA, LUAD, LUSC, KIRC, and HNSC) by keeping the samples which have both coding expression data and miRNA expression data for the five datasets respectively. We firstly take the intersection of the coding genes in a TCGA dataset with the genes in the PPI network (Vinayagam *et al.*, 2011) to get the list of coding genes with respect to that TCGA dataset. Then in the list of coding genes obtained, we use the TF list to distinguish TFs and mRNAs (i.e. the other coding genes excluding TF genes). We keep all obtained TFs since TFs play the key role in cell function (Vaquerizas *et al.*, 2009) and can be potential cancer drivers. However, for mRNAs and miRNAs, to reduce the number of genes, we firstly calculate the mean expression value for each gene and the standard deviation of each gene’s expression. Then we select the top 100 genes with the highest standard deviations and remove the other genes from the dataset. After the sample and gene selection, we have the expression data of 747 samples for BRCA, 445 samples for LUAD, 336 samples for LUSC, 240 samples for KIRC, and 485 samples for HNSC, for 100 miRNAs and 939 coding genes (including both TFs and mRNAs).

**Step 2.** Build miRNA-TF-mRNA network for each cancer patient. Following the idea of LIONESS (Kuijjer *et al.*, 2019) of building sample specific gene regulatory networks (details in Section 2.2.2), we build the miRNA-TF-mRNA network for each patient based on the matched miRNA, TF, and mRNA expression data. We firstly build a miRNA-TF-mRNA network using all patients’ expression data. Then for each patient, we build a miRNA-TF-mRNA network using the expression data of all patients excluding the current patient. In the two networks, nodes represent miRNAs/TFs/mRNAs. The edge weight is the Pearson correlation between the two nodes. Next, based on the two networks, we infer the network for the patient using the method in LIONESS.

**Step 3.** Refine patient-specific gene regulatory networks. For the refinement, we remove from each patient-specific network obtained in the previous Step the following: (1) the TF-TF, TF-mRNA, and mRNA-mRNA interactions which are not in the PPI network (Vinayagam *et al.*, 2011) since the interactions in this network are important for cellular information processing; (2) the miRNA-TF and miRNA-mRNA interactions which are not in TargetScan, miRTarBase, TarBase, or miRWalk; and (3) the TF-miRNA interactions which are not in TransmiR. Furthermore, to keep edges with strong interactions, we remove an edge from the patient-specific network if the absolute value of its weight is less than a threshold, which is the mean of the absolute weights of all edges in the network. Because the obtained networks are constructed from the gene expression data of a cancer type and gene interaction databases, they are more reliable and specific to the corresponding cancer type. In addition, although we use the gene interaction databases to refine the patient-specific gene networks, the personalised information of the networks may not be lost as the weights of the remained edges in each network are different. Then after refining the patient-specific networks by keeping edges with strong interactions, the networks are different and specific to each patient.

##### Stage (2) Identifying personalised cancer drivers

According to the Network Control method (Liu *et al.*, 2011) (details in Section 2.2.3), a network can be fully controlled by a minimum subset of nodes of the network, called Minimum Driver Node Set (MDNS). To capture the control machanism of the gene regulation, we apply the Network Control method to detect the MDNS of each sample-specific miRNA-TF-mRNA network constructed in Stage (1) above. Then we try to remove nodes of the network one by one and re-evaluate the MDNS. If the size of the MDNS increases, the removed node is a critical node in the network. In other words, when a critical node is removed from the network, a bigger MDNS is required to control the whole network. This indicates that critical nodes play the central controlling or driving role in the gene network and thus we consider them as candidate cancer drivers. Because these drivers are discovered based on the network for a specific patient, they are the predicted personalised cancer drivers for that patient.

#### 2.2.2 Constructing personalised gene regulation networks

*pDriver* applies LIONESS (Kuijjer *et al.*, 2019) to build gene regulatory networks for single patients. LIONESS considers that the network estimated from all samples is the average of the sample-specific networks which are estimated from individual samples. Based on this idea, LIONESS uses a linear framework to estimate sample-specific networks as follows.

We use the similar notation as that in Kuijjer *et al.* (2019). Let 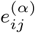 be the weight of the edge between node *i* and node *j* in the network obtained from all *N* samples (we denote the network as *α*). LIONESS assumes that 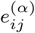 is the linear combination of the weights of the edge between nodes *i* and *j* in the *N* sample-specific networks:

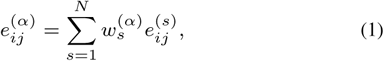

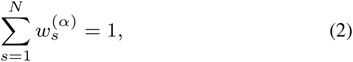

where 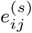 is the edge weight between node *i* and node *j* of sample (*s*), 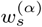 is the contribution of sample (*s*) to the aggregated network.

Similarly, the weight of the edge between node *i* and node *j* in the network from all samples except a sample *q* can be modelled as:

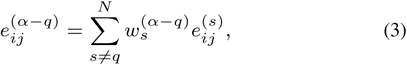

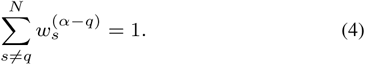

Suppose that the ratio of the contributions of a sample to the two networks (i.e. 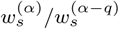) is the same for any sample *s ∈* {1, …, *N*} and *s ≠q*, from (2) and (4), we have:

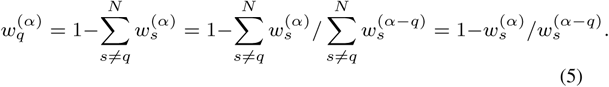

Subtracting Eq. 3 from Eq. 1, we find:

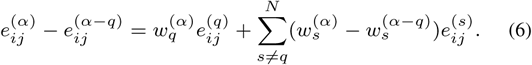

Replacing Eq. 6 with Eq. 5, we obtain:

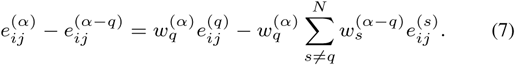

Thus, the weight of an edge of the network of sample *q* is computed:

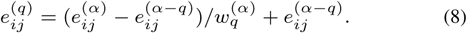

From Eq. 8, we can build the network for a specific sample based on the network from all samples and the network from all samples except the sample of interest, as illustrated in Fig. 1.

In *pDriver*, we assume that all samples make the same contribution to the network obtained from all *N* samples, which implies:

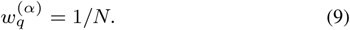

Replacing 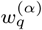 in Eq. 8 with 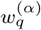 according to Eq. 9, we have:

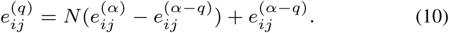

As we can see in Eq. 10, the mathematical framework is independent of the method used to compute the edge weights of the aggregated networks, we use Pearson, a common measure to quantify the association level between variables, to construct the network from all *N* samples and the network from *N* −1 samples. Since the output network from LIONESS contains all edges among nodes (i.e. a fully connected network), it may include untrue edges. To overcome this limitation, in *pDriver*, we refine the network obtained by LIONESS by using the existing databases, including protein protein interactions, miRNA-TF/mRNA interactions, and TF-miRNA interactions. Then we further refine the resulted network by removing edges which have the weight smaller than a threshold to assure the final network only contains reliable edges.

#### 2.2.3 Discovering personalised cancer drivers

Based on Kalman’s controllability rank condition (Kalman, 1963), the state and state transition of a system/network is fully controlled by a subset of nodes, but finding such a subset of nodes is computationally expensive or prohibitive for large networks like gene regulation networks. However, recently Liu *et al.* (2011) has provided an analytical method (called Network Control method) to detect the driver node set in a complex system modelled by a weighted directed network.

In this paper, we adopt this Network Control method to identify the driver node sets. Nevertheless, we only focus on the driver node set which has the smallest number of driver nodes, called the Minimum Driver Node Set (MDNS), a minimum subset of nodes which can fully control the network (Liu *et al.*, 2011) (The details are discussed in Section 1 of the Supplement). After we have the MDNS, we detect critical nodes of the network by removing nodes one by one out of the network. If the MDNS of the network with a node removed becomes larger, the removed node is a critical node. Since without the critical nodes, the size of the MDNS increases or we need to interact on more nodes to control the whole network, the critical nodes play the central role in controlling the network and they are considered as candidate cancer drivers.

#### 2.2.4 Implementation

The R source code of *pDriver* and the scripts to reproduce the experiment results in this paper are available at https://github.com/pvvhoang/pDriver.

## 3 Results

### 3.1 *pDriver* is robust in identifying coding cancer drivers

In this section, we compare the performance of *pDriver* with eight existing methods with different approaches for discovering cancer driver genes, including three methods for identifying personalised cancer drivers (DawnRank (Hou and Ma, 2014), PNC (Guo *et al.*, 2019), and SCS (Guo *et al.*, 2018)) and five methods for identifying cancer drivers at the population level (ActiveDriver (Reimand and Bader, 2013), DriverML (Han *et al.*, 2019), DriverNet (Bashashati *et al.*, 2012), MutSigCV (Lawrence *et al.*, 2013), and OncodriveFM (Gonzalez-Perez and Lopez-Bigas, 2012)). Besides, ActiveDriver, DriverML, MutSigCV, and OncodriveFM are mutation-based methods while DawnRank, DriverNet, PNC, and SCS are network-based methods. Since these methods are developed to identify only coding driver genes, we compare these methods with *pDriver* in unravelling coding driver genes.

As there is no ground truth for personalised drivers, following the same approach in existing methods for predicting personalised drivers, we aggregate the results of individuals to compare the performance of *pDriver* with the other methods. We firstly apply *pDriver* to detect patient-specific cancer driver genes. Based on the results, we compute the frequency of predicted cancer driver genes in the population. The more frequent a predicted cancer driver gene is, the higher it is in the ranking list of candidate cancer drivers at the population level.

From the five TCGA cancer datasets, BRCA, LUAD, LUSC, KIRC, and HNSC, we obtain the gene expression data of the respective cancer types. We use the results of the eight methods above for these selected cancer types in the PNC paper (Guo *et al.*, 2019) directly in the comparison.

We use CGC as the ground truth for predicted coding driver genes. CGC is from the COSMIC database (Forbes *et al.*, 2015) and it is a popular cancer gene dataset commonly used to validate cancer driver genes discovered by computational methods in cancer research. Furthermore, we use *F*_1_*Score* to measure the performance of the methods. *F*_1_*Score* assesses the enrichment ability of predicted cancer driver genes in the ground truth (i.e. CGC). *F*_1_*Score* is a combination of Precision and Recall, and it is computed as: 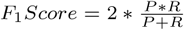, where *P* (Precision) indicates the fraction of correctly predicted driver genes among the predicted driver genes and *R* (Recall) represents the fraction of correctly predicted driver genes among the driver genes in the gold standard (i.e. CGC).

The results are shown in Fig. 2 (The *F*_1_*Scores* for all the methods are shown in Sections 2 of Supplementary materials and the lists of coding cancer drivers predicted by *pDriver* is shown in Sections 5 of Supplementary materials). It can be seen that the *F*_1_*Score* of *pDriver* is higher than those of the other eight methods, indicating the effectiveness of *pDriver* in identifying coding cancer drivers at the population level.

**Fig. 2.**
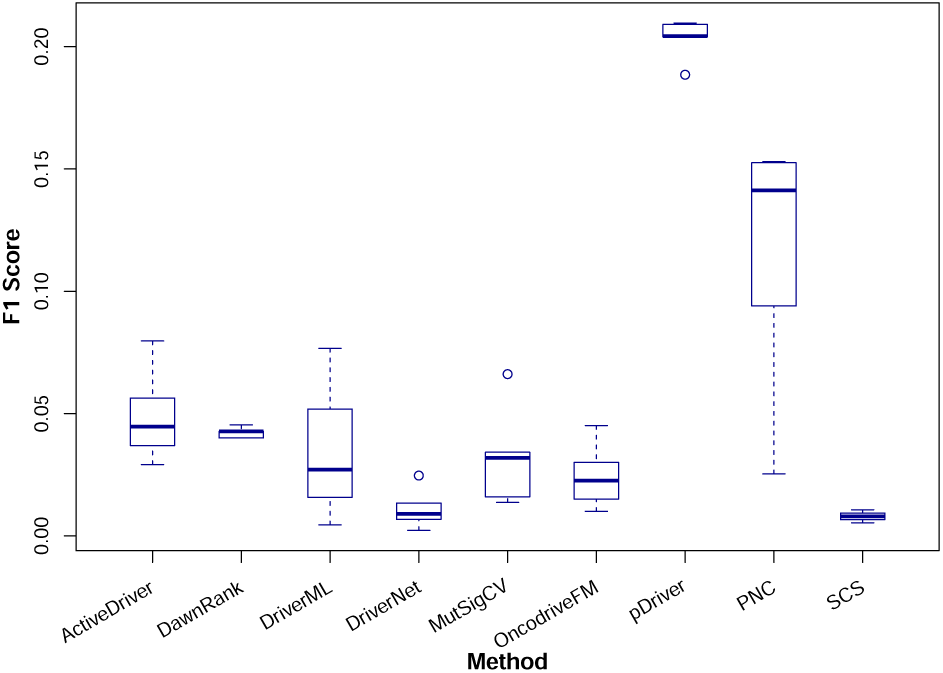
Comparison of F1 Score of the results by ActiveDriver, DawnRank, DriverML, DriverNet, MutSigCV, OncodriveFM, *pDriver*, PNC, and SCS. The x-axis shows the 9 methods and the y-axis is for *F*_1_*Score*. The results are computed based on 5 TCGA cancer datasets BRCA, LUAD, LUSC, KIRC, and HNSC.

Furthermore, to see if *pDriver* discovers similar cancer driver genes as the other methods, we compare the driver genes discovered by *pDriver* with those discovered by the top 5 performing methods (i.e. PNC, ActiveDriver, DawnRank, MutSigCV, and DriverML) among the eight. The findings are shown in Fig. 3. In the figure, the cancer drivers for the five cancer types (i.e. BRCA, LUAD, LUSC, KIRC, and HNSC) discovered by these methods are validated with the CGC and intersected to find the overlaps. Although there are some validated cancer drivers unravelled by different methods, *pDriver* uncovers a large amount of cancer driver genes which are not discovered by others. This shows that the results of *pDriver* and other methods are complementary, and they can be used together to improve the overall performance in cancer driver detection.

**Fig. 3.**
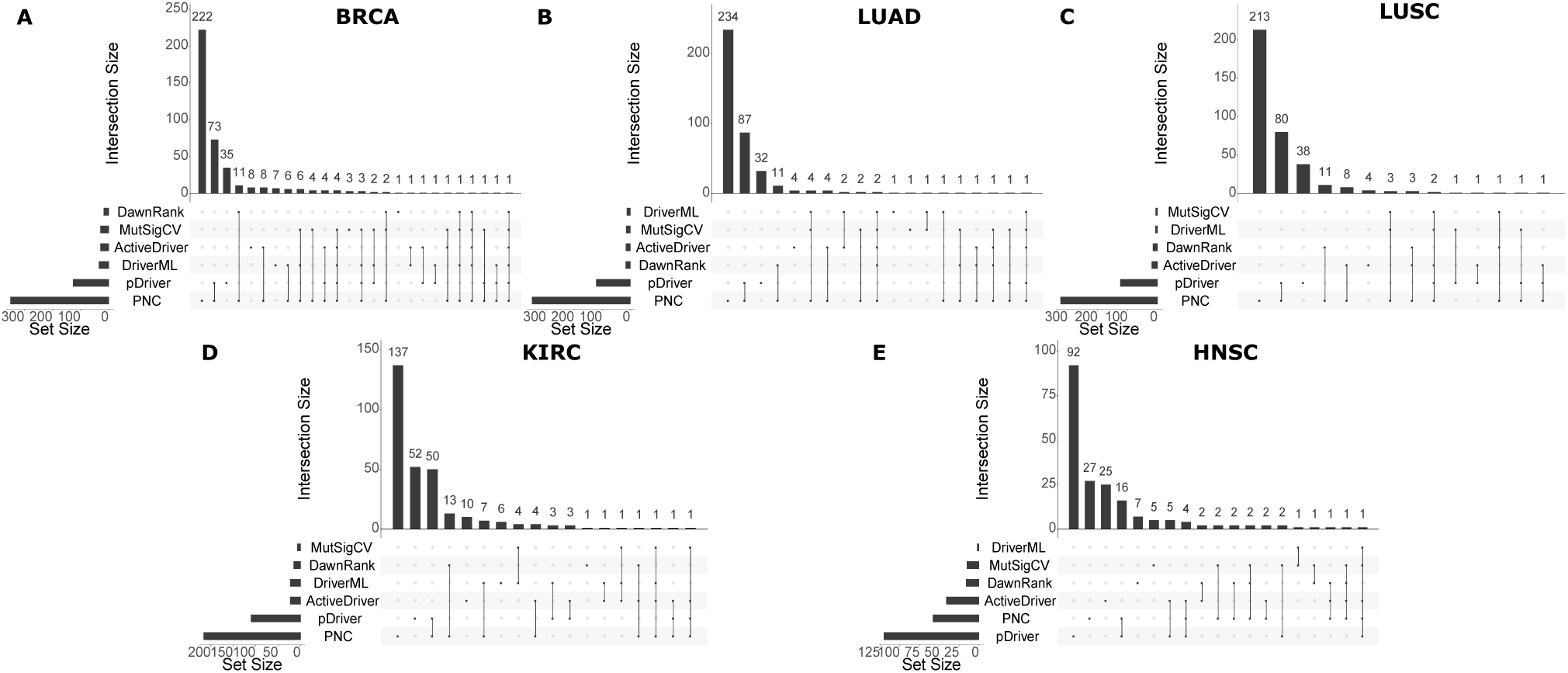
Overlap among different methods. The charts show the overlap between the cancer drivers discovered by the six methods (pDriver, PNC, ActiveDriver, DawnRank, MutSigCV, and DriverML) w.r.t each of the five cancer types (BRCA, LUAD, LUSC, KIRC, and HNSC). In each of the cases, the horizontal bars at the bottom left indicate the numbers of discovered cancer drivers which have been validated using the CGC, the vertical bars and the dotted lines together indicate the overlaps of validated cancer driver genes discovered by the methods.

### 3.2 Detecting miRNA cancer drivers

In addition to identifying coding cancer drivers, *pDriver* can also discover miRNA cancer drivers. As there is no ground truth for miRNA drivers, we use OncomiR (Wong *et al.*, 2018), a database of miRNA dysregulation in pan-cancer, to analyse the miRNA drivers predicted by *pDriver*. The percentages of the predicted miRNA drivers in OncomiR for the five cancer types (BRCA, LUAD, LUSC, KIRC, and HNSC) are shown in Fig. 4.

**Fig. 4.**
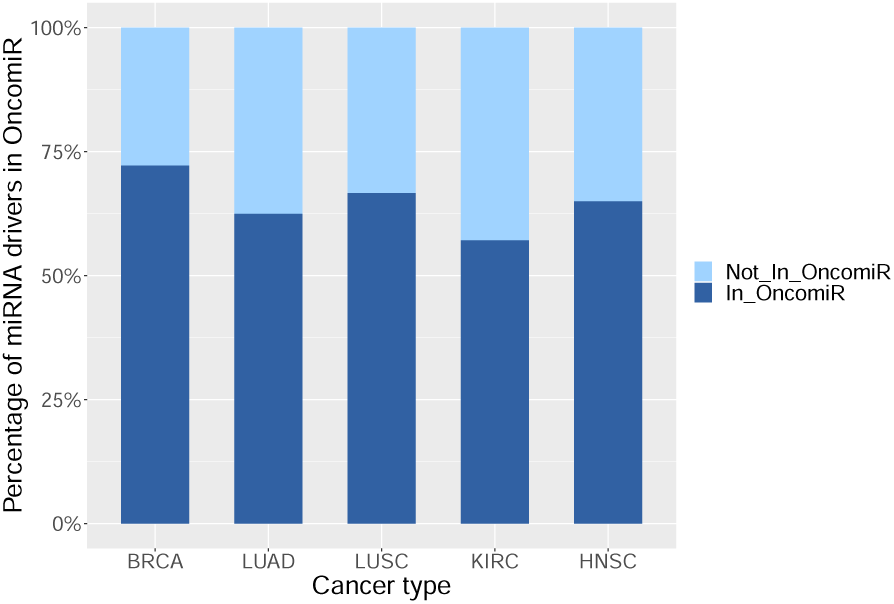
The percentage of the predicted miRNA drivers in OncomiR for BRCA, LUAD, LUSC, KIRC, and HNSC. The discovered miRNA cancer drivers at the population level are validated against OncomiR. Each bar indicates the percentage of miRNA drivers in OncomiR (dark blue) and not in OncomiR (light blue).

From Fig. 4, the percentages of predicted miRNA drivers for OncomiR are over 50% in all the five cancer types. Particularly, out of the 18 miRNAs which are identified by *pDriver* as BRCA drivers, 13 are recorded in OncomiR as involved in the turmorigenesis of BRCA. The p-value is 9.856e-06 based on the following hypergeometic test:

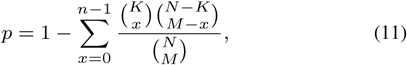

where *N* is the number of miRNAs of interest, *K* indicates the number of miRNAs in OncomiR, *M* denotes the number of predicted miRNA drivers, and *n* indicates the number of predicted miRNA drivers in OncomiR.

The figures for LUAD, LUSC, KIRC, and HNSC are 10/16, 14/21, 8/14, and 13/20 respectively. The corresponding p-values are at 7.783e-05, 7.192e-06, 0.002, and 4.890e-06.

Among the miRNA cancer drivers predicted by *pDriver*, there are many significant miRNAs which are confirmed to be related to different cancer types by other works. For example, *hsa-miR-375* in breast cancer cells experiences a high constitute expression (Frank *et al.*, 2019) and it is bound to estrogen receptor *α* to trigger off the transcription of the receptor in breast cancer (Yan *et al.*, 2014). The overexpression of *hsa-miR-940* in MDA-MB-231 breast cancer cells induces extensive osteoblastic lesions (Hashimoto *et al.*, 2018) and it is considered as a diagnostic and prognostic biomarker for breast cancer patients (Liu *et al.*, 2018). Other identified driver miRNAs involved in breast cancer include *hsa-miR-760* (Hu *et al.*, 2016), *hsa-miR-326* (Du *et al.*, 2019; Ghaemi *et al.*, 2019; Liang *et al.*, 2010), *hsa-miR-577* (Yin *et al.*, 2018), and *hsa-miR-429* (Pham *et al.*, 2019). In addition, *hsa-miR-326* also targets phoxa2 to regulate cell proliferation and migration in lung cancer (Wang *et al.*, 2016). *hsa-miR-1293, hsa-miR-1269a*, and *hsa-miR-1269b* are involved in the overall survival of kidney cancer patients (Liang *et al.*, 2017). Similarly, Luo *et al.* (2017) considers *hsa-miR-1293* as a prognostic biomarker for kidney cancer. The expression level of *hsa-miR-375* is used to predict head and neck cancer in the work of Avissar *et al.* (2009).

The list of miRNA drivers uncovered by *pDriver* at the population level for BRCA are in Table 1. The lists of miRNA drivers for other cancer types are presented in Section 6 of the Supplement.

**Table 1.**
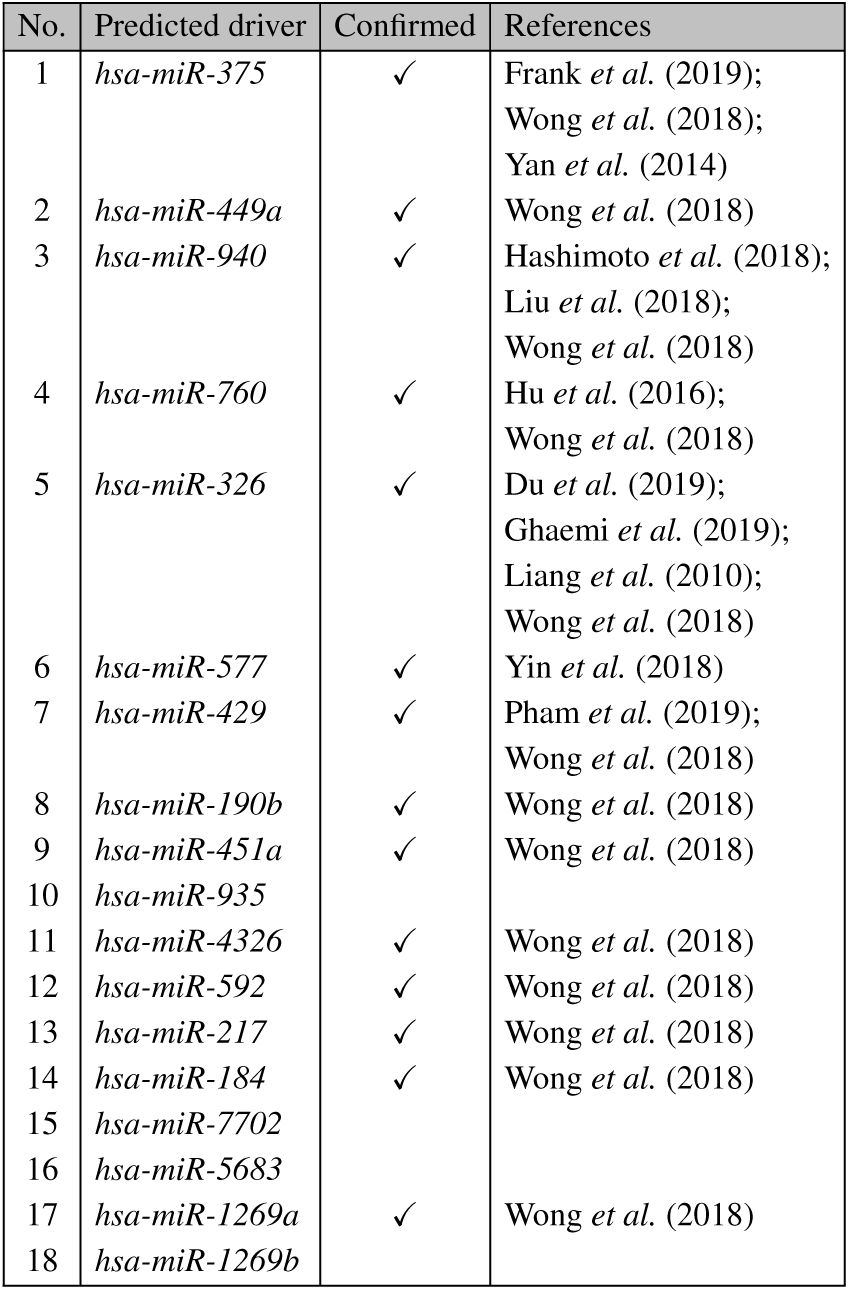
Predicted miRNA drivers in BRCA at the population level

### 3.3 Discovering personalised cancer drivers

In this section, we discuss the ability of *pDriver* to discover personalised coding and miRNA cancer drivers, which differentiates *pDriver* from other existing methods. On average, *pDriver* identifies 99.98 coding and miRNA drivers for an BRCA individual patient. The figures for LUAD, LUSC, KIRC, and HNSC are 94.95, 98.55, 98.68, and 103.85 respectively.

For personalised coding cancer drivers, we focus on rare drivers (i.e. drivers are mutated in a small number of patients) since these driver genes have a low mutation frequency, they are usually missed by other methods, especially the methods based on the mutation information. In our case study, we consider rare drivers are mutated in less than 0.5% of patients but they are predicted as personalised cancer drivers in different patients by *pDriver*. Taking the breast cancer as example and the mutation data from TCGA, we obtain 58 personalised rare coding drivers (see the list of these drivers in Section 7 of the Supplement).

We use Enrichr (Kuleshov *et al.*, 2016) to analyse the functional enrichment of these 58 drivers. Table 2 shows that these driver genes are significantly enriched (adjusted p-value < 0.05) in various GO biological processes and KEGG pathways associated with breast cancer, such as cell differentiation (GO:0045597, GO:0045598, GO:1902036, etc.) (Jogi *et al.*, 2012), cell proliferation (GO:0008285, GO:2000178, GO:2000177) (Ciocca and Fanelli, 1997), signaling pathway (Nwabo Kamdje *et al.*, 2014). These results indicate that the predicted rare coding drivers are closely related to the biological condition of breast cancer and they are biologically meaningful.

**Table 2.** Enriched GO biological processes and KEGG pathways related to breast cancer of the predicted rare coding drivers

| Term | #Genes | p-value |
| --- | --- | --- |
| Ontologies |  |  |
| GO:0045597 positive regulation of cell differentiation | 6 | 2.6e-03 |
| GO:0045598 regulation of fat cell differentiation | 4 | 5.7e-03 |
| GO:1902036 regulation of hematopoietic stem cell differentiation | 4 | 6.9e-03 |
| GO:1901532 regulation of hematopoietic progenitor cell differentiation | 4 | 7.1e-03 |
| GO:1901532 regulation of hematopoietic progenitor cell differentiation | 4 | 7.1e-03 |
| GO:0045596 negative regulation of cell differentiation | 5 | 7.0e-03 |
| GO:0045639 positive regulation of myeloid cell differentiation | 3 | 7.4e-03 |
| GO:0008285 negative regulation of cell proliferation | 7 | 7.6e-03 |
| GO:0045599 negative regulation of fat cell differentiation | 3 | 9.4e-03 |
| GO:2000178 negative regulation of neural precursor cell proliferation | 2 | 1.0e-02 |
| GO:2000736 regulation of stem cell differentiation | 4 | 1.0e-02 |
| GO:2001014 regulation of skeletal muscle cell differentiation | 2 | 2.0e-02 |
| GO:0045595 regulation of cell differentiation | 4 | 2.7e-02 |
| GO:2000177 regulation of neural precursor cell proliferation | 2 | 4.2e-02 |
| Pathways |  |  |
| TGF-beta signaling pathway | 8 | 1.2e-08 |
| Hippo signaling pathway | 6 | 1.9e-04 |
| Thyroid hormone signaling pathway | 5 | 4.4e-04 |
| Relaxin signaling pathway | 5 | 6.8e-04 |
| Adipocytokine signaling pathway | 4 | 7.8e-04 |
| B cell receptor signaling pathway | 4 | 8.2e-04 |
| PPAR signaling pathway | 4 | 9.2e-04 |
| MAPK signaling pathway | 6 | 2.5e-03 |
| T cell receptor signaling pathway | 4 | 2.4e-03 |
| C-type lectin receptor signaling pathway | 4 | 2.5e-03 |
| TNF signaling pathway | 4 | 3.0e-03 |
| Estrogen signaling pathway | 4 | 6.5e-03 |
| Wnt signaling pathway | 4 | 8.6e-03 |
| cGMP-PKG signaling pathway | 4 | 9.8e-03 |
| IL-17 signaling pathway | 3 | 1.6e-02 |
| AGE-RAGE signaling pathway in diabetic complications | 3 | 1.9e-02 |
| cAMP signaling pathway | 4 | 2.0e-02 |
| Toll-like receptor signaling pathway | 3 | 2.0e-02 |
| Oxytocin signaling pathway | 3 | 4.9e-02 |

Among the predicted rare coding drivers, there are some significant genes such as *JUN, CREB1*, and *ID2*, which are enriched in numerous biological processes and pathways. Particularly, *JUN* and *CREB1* are involved in the process of positive regulation of cell differentiation (GO:0045597), TNF signaling pathway, cAMP signaling pathway, etc. while *ID2* is associated with the process of negative regulation of cell proliferation (GO:0008285), TGF-beta signaling pathway, hippo signaling pathway, etc. Although these three genes have a low mutation frequency in breast cancer patients, they play a vital role in the gene network of breast cancer patient TCGA-AC-A62Y and they are predicted as cancer drivers of this patient. We further analyse all predicted cancer drivers of the breast cancer patient TCGA-AC-A62Y and we see that this patient has 91 coding cancer drivers with 37 driver genes in CGC. Especially, 11 of the 91 predicted driver genes (including *NCOA1, JUN, SP1, MYC, NCOA3, LEF1, PGR, E2F3, FOS, TP53*, and *ESR1*) are enriched in the KEGG breast cancer pathway (adjusted p-value = 2.3e-09). Other driver genes are also significantly enriched in several biological processes and pathways related to breast cancer such as regulation of fat cell proliferation (GO:0070344), TGF-beta signaling pathway, estrogen signaling pathway, etc.

In addition, the patient TCGA-AC-A62Y has 11 miRNA cancer drivers discovered by *pDriver*. Among these 11 miRNA drivers, 9 miRNAs are confirmed to be associated with breast cancer by OncomiR (Wong *et al.*, 2018). Furthermore, 4 miRNAs (including *hsa-miR-449a, hsa-miR-375, hsa-miR-451a*, and *hsa-miR-326*) are enriched in the KEGG pathway, named microRNAs in cancer, with the adjusted p-value being 0.005. *hsa-miR-577* targets *RAB25* to suppress epithelial-mesenchymal transition and metastasis of breast cancer (Yin *et al.*, 2018). The detected miRNA driver *hsa-miR-935* is novel and it can be used as a candidate cancer driver in web-lab experiments to confirm its role in the cancer development of the patient TCGA-AC-A62Y.

### 3.4 Distribution of personalised rare cancer drivers in breast cancer subtypes

Breast cancer includes numerous subtypes and each subtype has specific morphologies as well as clinical outcomes. Thus, to elucidate the difference of breast cancer subtypes, we analyse the distribution of personalised rare cancer drivers in patients across the four major breast cancer subtypes, including Basal, Her2, Luminal A (LumA), and Luminal B (LumB).

In Section 3.3, we use 0.5% as a cutoff of mutation frequency to determine rare drivers. However, to better visualise the distribution of personalised rare drivers among breast cancer subtypes, we use different cutoffs (i.e. 0.2%, 0.3%, 0.4%, and 0.5%) to obtain rare drivers in this section. The rare drivers identified by these cutoffs are not cumulative. Then we validate these rare drivers with CGC and the numbers of validated rare drivers are 5, 4, 3, and 6 accordingly.

To identify the subtype of a breast cancer patient, we apply the Pam50 method (Liu *et al.*, 2016; Parker *et al.*, 2009) for the breast cancer dataset. We obtain 158 patients in Basal subtype, 108 patients in Her2, 221 patients in LumA, 165 patients in LumB, and 95 patients in Normal-like. In this case study, we only focus on the patients in the four major subtypes of breast cancer (i.e. Basal, Her2, LumA, and LumB).

We then compute the proportion of patients in each breast cancer subtype, who are driven by all the validated rare drivers according to each mutation frequency cutoff. The distribution results are shown in Fig. 5. As can be seen from the figure, the distribution of rare drivers in Basal patients is different from the distributions of patients in other subtypes. Basal patients tend to have the least common rare drivers compared to other subtypes. In other words, they are driven by different rare drivers. This result might be related to the great heterogeneity of Basal breast cancer. According to Cheang *et al.* (2008), Basal patients are difficult to be treated because of their tumour heterogeneity and they have a poor prognosis. On the other hand, other cancer subtype patients such as LumA have a less heterogeneity and they have a good prognosis. When LumA patients are treated early, they usually have the most favourable outcome and experience a lower chance of relapse (Guedj *et al.*, 2012; Neve *et al.*, 2006).

**Fig. 5.**
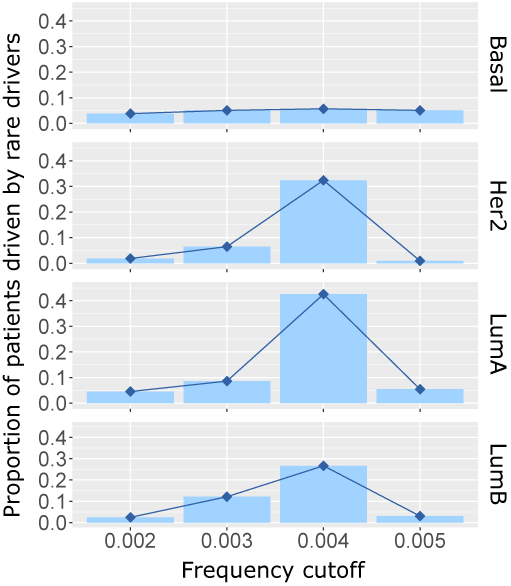
The distribution of the rare coding drivers in CGC across breast cancer subtypes (Basal, Her2, LumA, and LumB). The x-axis indicates mutation frequency cutoffs from 0.2% to 0.5% and these cutoffs are not cumulative. The y-axis shows the proportion of patients driven by the rare drivers within that cutoff in the corresponding cancer subtype.

### 3.5 The effectiveness of the Network Control method for finding influential nodes in a network

To assess the effectiveness of *pDriver* applying the Network Control method in identifying driver genes in the personalised networks, we compare the performance of the Network Control method with the other two methods, PageRank (Brin and Page, 1998; Page *et al.*, 1998) and Influence Maximisation (IM) (Gong *et al.*, 2016; Yang *et al.*, 2016b) when use each of them in Stage (2) of *pDriver* (the details of PageRank and IM are discussed in Section 3 and 4 of the Supplement respectively).

PageRank is used by Google to rank the importance of webpages. In this case study, we apply PageRank to identify important genes in biological networks, similar to searching for important webpages over the Internet. IM is usually used to discover users who have high influence on other users in online social networks. We apply IM to detect genes which have great influence on other genes in the miRNA-TF-mRNA networks. IM identifies an influential node (assuming it is active) by evaluating the amount of other nodes which can be activated by this active node. The more nodes the node can activate, the greater influence it has in the network. To evaluate the influence of nodes, IM requires the information of node weights, which are used to determine if an inactive node can be activated or not. We set the weight of a node (gene) as the absolute difference between average expression of the gene in normal state and its average expression in tumour state. An inactive node can be activated if the total weights of the edges from its active neighbouring nodes to it is larger than its weight. Intuitively, if the total effect from active neighbouring nodes on an inactive node is strong enough, they can change the state of that node from inactive to active. In our case study, we consider inactive and active states as normal and tumour states respectively. As the node weight is the expression difference of a gene between normal and tumour state, it is the cost required to change that gene from normal state to tumour state. If a gene has a large weight, it requires a high cost to change between its states.

We use the BRCA dataset to evaluate these methods in identifying driver genes in patient-specific networks. We use the same patient-specific networks built from Stage (1) of *pDriver* for all the methods. In addition, we use the same method to aggregate the results of individual results as described in Section 3.1. We also use CGC as the gold standard. We assess the performance of these methods based on the *F*_1_*Scores* of coding driver discovery at the population level.

We choose the top 100, 200, 300, and 400 discovered cancer drivers by using each of the three methods and validate with CGC. The comparison result is shown in Fig. 6. In all the cases, the performance of the Network Control method is comparable to or better than the other two methods. It indicates that the Network Control method is more effective than other methods in evaluating the role of genes in biological networks. The reason for this result may come from the fact that gene regulation is related to the control mechanism, which has been captured by the Network Control method while the other two methods discover influential nodes from the perspective of information propagation. Thus, the Network Control method is a better choice comparing to other two network based methods for finding influential nodes in a network.

**Fig. 6.**
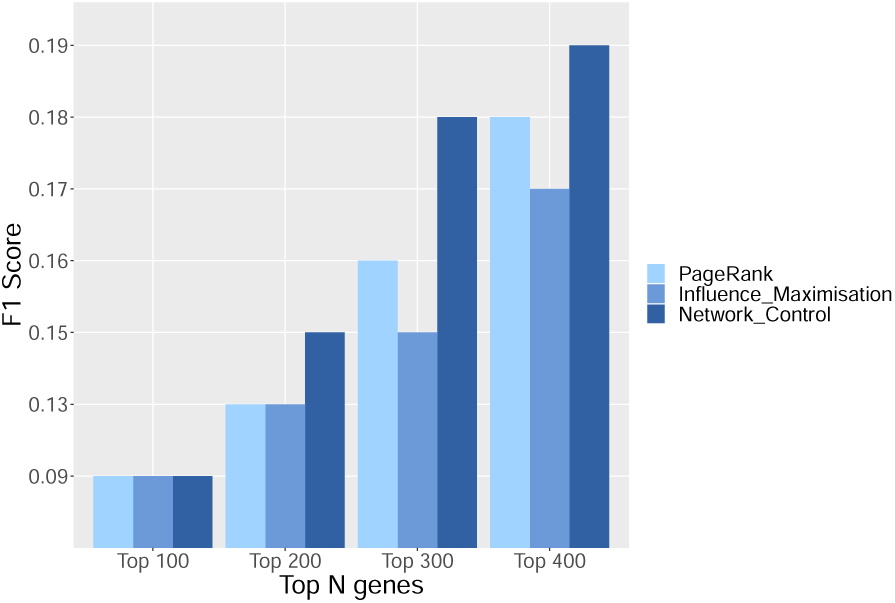
Comparison of the effectiveness of PageRank, Influence Maximisation, and Network Control method in identifying influential genes in the gene networks. The discovered coding cancer drivers at the population level using the three methods are validated against CGC. Each bar indicates the *F*_1_*Score* of driver gene prediction with each method according to the top 100, 200, 300, and 400 discovered driver genes.

## 4 Conclusion

Because each cancer patient possesses a different genome, their disease may be driven by different cancer driver genes. As a result, two cancer patients who have the same cancer type and experience the same treatment may have different outcomes. However, most of the current methods discover cancer drivers at the population level. Recently some methods have been developed to detect personalised cancer drivers, but they do not use the gene network information specific to each patient, or they require the gene expression data of a tumour and its neighbour, which is not always available. In addition, these methods only uncover coding drivers while non-coding genes also play an important role in cancer development. Thus, it is necessary to develop novel methods to identify personalised cancer drivers, including both coding and non-coding drivers, to elucidate their regulatory mechanism in cancer patients.

To overcome the limitations, we have proposed the novel method, *pDriver*, to unravel personalised coding and miRNA cancer drivers by using the gene expression data and the target binding information of genes. We have assessed the performance of *pDriver* with different experiments. The results demonstrate that *pDriver* is more effective than other existing methods in identifying population-level coding cancer drivers. *pDriver* can discover miRNA cancer drivers and most of them are confirmed to be involved in cancer by literature. *pDriver* can also detect personalised rare coding drivers. These predicted drivers are significantly enriched in various GO biological processes and KEGG pathways related to cancer. Furthermore, we have found that Basal cancer patients have different personalised rare coding drivers, which may explain the greater heterogeneity of Basal breast cancer. In summary, all the results indicate that *pDriver* can provide new insights into the molecular regulatory mechanisms in the development of cancer and it has the potential to contribute significantly to the design of effective cancer treatments.

## Acknowledgements

This research is supported by the Australian Government Research Training Program (RTP) Scholarship and the Vice Chancellor & President’s Scholarship offered by the University of South Australia.

